# Amenability to Engineering of the Homologation Enzyme, HphA, through Homologous-Based Site-Directed Mutagenesis

**DOI:** 10.64898/2025.12.01.691582

**Authors:** Rebecca M. Lang Harman, Gracie Blackstone, Favor O. Aruna, Shivam R. Patel, Minho Shin, Reed K. NeSmith, D. Brooks Dickson, Angela C. Spencer, Shogo Mori

## Abstract

Homologation of amino acids, the addition or deletion of a methylene group onto their side chains, has the potential to increase the biostability and bioavailability of peptide natural products. The first enzyme in the homologation pathway, HphA, has been previously characterized and is substrate selective. Bioinformatics studies were used to identify amino acids in the active site of HphA, which may play a role in substrate selection, by comparison to homologous enzymes, homocitrate synthase (HCS) and 2-isopropylmalate synthase (IPMS). Single point mutants to five amino acid residues in the HphA’s active site were created to mimic those of HCS and IPMS. Their activities were measured via time-course assays with the natural substrates for HCS and IPMS. Residue A73 was identified as important in the substrate specificity of HphA; therefore, six different additional mutations were generated and tested with nine substrates with various side chains. The HphA A73L mutant exhibited the highest activity compared to the other mutants, showing activity with counterparts of L-Tyr (HphA natural substrate), L-Val (IPMS natural substrate), L-Leu, L-Ser, L-Trp, and L-Asp. Kinetic assays were taken with HphA A73L with the active substrates and compared with kinetic data from HphA WT, HCS, and IPMS. These results demonstrated that the A73L mutation significantly relaxed the substrate specificity of HphA, indicating its amenability to engineering. This research will serve as the foundation for future metabolic engineering studies on the enzymatic homologation pathway of amino acids.

## INTRODUCTION

Natural products (NPs) are compounds found in nature that are synthesized by living organisms.^1^ These substances are characterized by their high molecular weight and structural complexity, properties which may be difficult to achieve via traditional organic chemistry methods.^*2*^ These features make NPs an essential component in the development of small molecule drugs.^3^ The medicinally useful NPs include peptide NPs due to their high diversity and biological activities.^4^ Most of them are categorized as either nonribosomal peptides (NRPs) or ribosomally synthesized and post-translationally modified peptides (RiPPs). RiPPs are synthesized using only canonical amino acids but are extensively post-translationally modified to produce diverse structures.^5^ NRPs exhibit significant structural diversity, with many containing non-proteinogenic amino acids, which enhance the biostability and bioavailability of peptide natural products.^6^

Homologation of amino acids is the insertion or deletion of a methylene group in the side chain of an amino acid (Figure 1), generating a non-proteinogenic amino acid in a peptide NP. It is hypothesized that this modification increases the stability and bioavailability of the peptide NP.^7^ One of the common NP families that contain homologated amino acids (homoAAs) is anabaenopeptins produced by a variety of cyanobacteria.^8^ The genes responsible for the homologation of L-Phe and L-Tyr in anabaenopeptin biosynthesis in the cyanobacterium *Nostoc punctiforme* PCC 73102, *hphA/B/CD*, were previously identified, also noting the high substrate selectivity of the homologation pathway.^9^ Previous metabolic engineering studies on the homologous pathways in the bacterial and plant primary metabolism for L-Val^10^ and L-Met^11^ homologations, respectively, suggested that the first enzyme in the pathway, HphA, acts as the gatekeeper. Indeed, HphA was recently characterized as a highly selective enzyme.^12^ While modifying existing NPs using known homologation pathways would be advantageous, the high substrate selectivity of HphA remains a significant obstacle.

**Figure 1.**
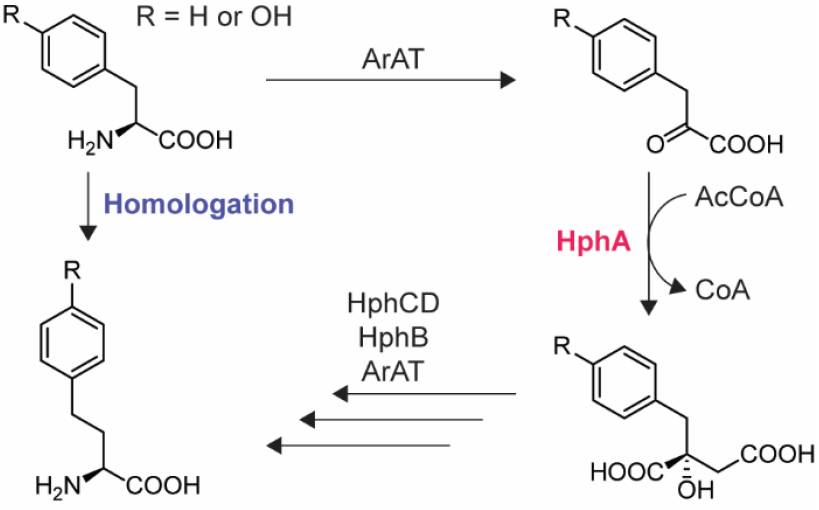
Homologation pathway for L-Phe and L-Tyr, focusing on the target enzyme HphA in this study. Each abbreviation depicts as follows: ArAT = aromatic amino acid aminotransferase, AcCoA = acetyl coenzyme A, CoA = coenzyme A.

The goal of this study was to determine whether HphA is amenable to engineering. Bioinformatics was used to design mutants to mimic the activity of its homologous enzymes, homocitrate synthase (HCS) and 2-isopropylmalate synthase (IPMS) for L-Lys and L-Leu biosynthesis, respectively. Site-directed mutagenesis was used to insert the mutations, and the activity of the mutant enzymes was characterized using colorimetric assays with Ellman’s reagent.^13^ The amino acid residue in the HphA active site that is responsible for the selection of the substrate was identified; the amenability of HphA to engineering was demonstrated by relaxing its substrate selectivity.

## MATERIALS AND METHODS

### Bacterial Strains, Plasmids, Materials, and Instrumentation

Chemically competent *Escherichia coli* DH5α and BL21(DE3) cells were purchased from Thermo Scientific (Waltham, MA) and Invitrogen (Waltham, MA), respectively. DNA oligonucleotides were purchased from Integrated DNA Technologies (Coralville, IA). The pET-28a overexpression plasmid for *E. coli* was purchased from Novagen (MilliporeSigma, Burlington, MA). DNA sequencing was performed by Eurofins Genomics USA (Louisville, KY). Small molecule chemicals and bacterial media were purchased from Fisher Scientific (Waltham, MA), MilliporeSigma, AA Blocks (San Diego, CA), and CoALA Biosciences (Elgin, TX) and used without modifications. Enzymes and buffers for the molecular biology experiments were purchased from New England Biolabs (Ipswich, MA) and used as instructed.

### Bioinformatics analysis of HphA, IPMS, HCS, and hHphA

The amino acid sequence alignments between HphA (NCBI accession number: WP_012409019) and HCS as well as HphA and IPMS were performed in the previous study.^12^ The amino acid sequence of HphA from *Microcystis aeruginosa* NIES-4285 (hHphA) was obtained from the National Center for Biotechnology Information (NCBI; accession number: WP_130757515).^14^ The MultAlin tool (http://multalin.toulouse.inra.fr/multalin/) was used to align the amino acid sequences.^15^

### Site-directed mutagenesis of HphA

Site-directed mutagenesis of HphA was conducted using overlap extension polymerase chain reaction (PCR).^16^ All PCRs were performed with Phusion DNA polymerase (New England Biolabs) using the following components: Phusion GC buffer, 0.2 mM dNTP, 20 pg/µL template, 0.5 mM forward and reverse primers, 3% dimethyl sulfoxide (DMSO), Phusion DNA polymerase, and water to a total volume of 50 µL. The PCR programs were optimized as follows: initial denaturation for 1 min at 95 °C, 30 cycles of 20 s at 95 °C, 30 s at 52 °C, and 2 min and 30 s at 72 °C, and a final extension for 5 min at 72 °C. The template for the site-directed mutagenesis was the HphA wild-type (WT) expression plasmid, pHphA-CHis-pET28a, which was constructed in a previous study.^12^ All HphA mutant expression plasmids were constructed so that the expressed protein contains a 6×His tag at the C-terminus. The mutagenesis began with synthesizing two DNA fragments that contain the corresponding mutations in their overlapped regions through PCR. This first round of PCR was performed using two primer pairs: (1) primer #1 with the reverse mutation primer (HphA-Mutation-R) and (2) primer #2 with the forward mutation primer (HphA-Mutation-F) (Table S1). These two PCR products were subsequently purified through agarose gel extraction (GeneJET, Thermo Scientific) and served as templates for the second round of PCR. In the second round of PCR, the primer pair consisting of primer #1 and primer #2 was employed to generate the full-length *hphA* mutant. This PCR product was also purified via agarose gel extraction and digested by the restriction enzymes NcoI and XhoI, along with the pET-28a plasmid. The digested *hphA* mutant and pET-28a were ligated with T4 DNA ligase, and the ligation product was then transformed into chemically competent *E. coli* DH5α. Each transformed single colony was cultured overnight to miniprep the recombinant plasmid, which was verified through double digestion followed by sequencing reactions.

### Overexpression and purification of HphA mutants

HphA mutants were overexpressed and purified for *in vitro* characterization. The methods for expression and purification of HphA mutants followed those used for HphA WT developed in a previous study.^12^ The recombinant expression plasmid was transformed into *E. coli* BL21(DE3) cells. Three colonies were picked up and cultured in 3 mL LB medium containing 50 µg/mL kanamycin at 37 °C with shaking at 200 rpm until the culture reached the log phase and appeared cloudy. This culture was then inoculated into 1 L of LB medium with 50 µg/mL kanamycin and shaken under the same conditions until the optimal density at 600 nm (OD_600_) reached approximately 0.5. After cooling the culture below 16 °C, 0.2 mM of isopropyl 1-thio-β-D-galactopyranoside (IPTG) was added, and the culture was incubated overnight at 16 °C with shaking at 200 rpm. The cells were harvested by centrifugation at 4,000 rpm for 15 min at 4 °C, which was then resuspended in approximately 30 mL of the Ni-NTA binding buffer (25 mM Tris-HCl pH 8.0, 400 mM NaCl, 5 mM imidazole, and 10% glycerol). Four cycles of sonication, each lasting 120 s with alternating 10 s “on” and 10 s “off,” was utilized to disrupt the cells. Cell debris was removed by centrifugation at 14,000 rpm for 45 min at 4 °C. The supernatant containing the solubilized proteins was incubated with Ni-NTA resin (Thermo Scientific) for 2 h at 4 °C with gentle rotation. The Ni-NTA resin was packed into a column and washed 10 times with 5 mL of the wash buffer (25 mM Tris-HCl pH 8.0, 400 mM NaCl, 40 mM imidazole, 10% glycerol). The protein was eluted in three fractions, using 1 mL of the elution buffer (25 mM Tris-HCl pH 8.0, 400 mM NaCl, 250 mM imidazole, and 10% glycerol) for each fraction. The first two elution fractions were dialyzed against a dialysis buffer (40 mM Tris-HCl pH 8.0, 200 mM NaCl, 2 mM β-mercaptoethanol, and 10% glycerol). Three dialysis solutions were used, with each dialysis session lasting at least three hours in a 1 L dialysis buffer. The protein concentration was measured using a spectrophotometer at λ = 280 nm. Finally, the protein solution was flash frozen in liquid nitrogen and stored at −80 °C.

### In vitro colorimetric assays for HphA mutants

Colorimetric assays for the HphA mutants were performed using Ellman’s reagent (5,5’-dithiobis-(2-nitrobenzoic acid) (DTNB))^13^ to measure product formation through the release of CoA following the enzymatic reaction. The reactions (100 µL) were prepared to a final concentration of 50 mM Tris-HCl (pH 7.5), 20 mM MgCl_2_, 20 mM KCl, 1 mM DTNB, 1.5 mM acetyl coenzyme A (AcCoA), 5 µM of enzyme, and 1 mM of substrate; these methods were previously developed for characterizing the wild-type HphA^14^ from information gathered on testing the activity of IPMS.^17^ Substrates were added directly to the 96-well plate, and the rest of the reaction mixture was prepared separately and then added to the corresponding well to start the reaction, after which the plate was put directly into the plate reader (xMark, Bio-Rad Laboratories, Hercules, CA). Reactions were measured at 412 nm every 15 seconds with orbital mixing for 1 s in between measurements. All colorimetric assays were duplicated.

### Substrate profile establishment for HphA mutants

Substrate profiling of HphA mutants was established using the same reaction and measurement conditions as mentioned in *In vitro colorimetric assays for HphA mutants*, with reactions run for 20 min each. To establish the activity of HphA single mutants, 3-methyl-2-oxobutanoic acid (MOBA) was used as a substrate for the IPMS mimics, alpha-ketoglutaric acid (αKG) and oxalacetic acid (OAA) were used for the HCS mimics, and homo-phenylpyruvic acid (hPPA) was used for the hHphA mimic. For all the A73 single mutants, the substrates were 4-hydroxylphenylpyruvic acid (4HPPA), MOBA, 4-methyl-2-oxovaleric acid (MOVA), aKG, OAA, hPPA, indole-3-pyruvic acid (InPA), imidazolepyruvic acid (ImPA), and hydroxypyruvic acid (HPA), with DMSO as a blank. The ImPA substrate was in hydrobromide hydrate form (C_6_H_6_N_2_O_3_•xHBr•yH_2_O), so its weight was calculated assuming x and y both equaled one, which meant that its formal weight was 253.05 g/mol. Both HphA A73G and A73D were not stable in the reaction solution, so their reaction solutions were prepared with 10% glycerol. To prevent the heat shock for these mutants, substrates were added to the microfuge tubes on ice to start the reactions and then immediately placed into the well plate for the measurement. All reactions were duplicated. Percent activities of the mutants were calculated by averaging the absorbance differences from 0 to 20 minutes of each reaction. The activities were then converted to percentage form, with HphA WT with 4HPPA activity being set to 100%, and all other mutants were related to that number.

### Kinetic assays for HphA

The colorimetric assays to calculate the Michaelis-Menten kinetic parameters for each active mutant were established by using the same conditions as previously mentioned (*In vitro colorimetric assays for HphA mutants*). The kinetic assays were run for HphA A73L with 4HPPA, MOBA, MOVA, OAA, InPA, and HPA. The assays were also run for HphA WT with MOBA and MOVA, IPMS with MOBA and MOVA, and HCS with aKG and OAA. Substrate concentrations varied depending upon the enzyme and substrate pair, but these ranged from 20 mM to 0.001 mM. For HphA WT, MOBA concentrations were 0.05, 0.1, 0.2, 0.5, 1, 2, 4.2, 5, 8.4, and 20 mM and MOVA concentrations were 0.05, 0.2, 0.5, 1, 3, and 5 mM. For IPMS, MOBA concentrations were 0.02, 0.05, 0.2, 0.5, 1, and 3 mM and MOVA concentrations were 0.002, 0.005, 0.02, 0.05, 0.2, and 0.5 mM. For HCS, aKG concentrations were 0.02, 0.05, 0.2, 0.5, 1, and 2 mM and OAA concentrations were 0.2, 0.5, 1, and 5 mM. HCS was not stable in the reaction solution, so the reactions were conducted with 10% glycerol and the substrate added on ice. For HphA A73L, 4HPPA concentrations were 0.01, 0.05, 0.1, 0.5, 1, and 3 mM; MOBA concentrations were 0.01, 0.05, 0.1, 0.5, 1, and 2 mM; MOVA concentrations were 0.01, 0.05, 0.1, 0.5, 1, 2, and 5 mM; OAA concentrations were 0.001, 0.01, 0.05, 0.1, 0.5, and 1 mM; InPA concentrations were 0.005, 0.01, 0.05, 0.1, 0.5, 1, 2, and 3 mM; and HPA concentrations were 0.02, 0.05, 0.2, 0.5, 1, and 2 mM.

After all the data was obtained, data of colorimetric assays run with each substrate with dialysis buffer as a negative control (*Overexpression and purification of HphA mutants*) was then subtracted from all points. Then, absorbance was converted to concentration utilizing a standard curve for thiol generated in-house using cysteine in the reaction solution. The calculated extinction coefficient was 3.8663 mM^-1^. The linear portions of the two reactions for each substrate concentration were taken and analyzed with GraphPad Prism (GraphPad Software, Boston, MA) to obtain Michaelis-Menten curves and parameters for each enzyme and substrate pair.

## RESULTS AND DISCUSSION

### Bioinformatics to design the mutant that mimics the activity of homologous enzymes

To show the amenability of HphA to engineering, bioinformatics was used to design mutants of HphA to mimic the activity of homologous enzymes in primary metabolisms. The homologous enzymes to HphA are HCS, IPMS (LeuA), and homoHphA (hHphA). HCS is the first enzyme involved in L-Lys biosynthesis after transamination, which is a regulated step that involves the condensation of AcCoA and αKG to form homocitrate and CoA.^18, 19^ IPMS is the first enzyme involved in L-Leu biosynthesis, and it catalyzes the condensation of MOBA and AcCoA into 2-isopropylmalate and CoA.^20^ hHphA is hypothesized to be involved in the production of doubly (di)-homologated L-tyrosine (di-hTyr).^14^

The bioinformatics analysis was performed in a previous study, which identified potential amino acid residues that are crucial for the selection of the substrate in HphA, HCSs, and IPMSs.^12^ It was first conducted with the most homologous structurally characterized enzyme, HCS from *Sulfolobus acidocaldarious* (PDB ID: 6KTQ, 28.9% sequence identity, 48.9% similarity to HphA),^19^ by taking advantage of this crystal structure binding the substrate αKG (Figure 2A). The amino acid sequence of another HCS from *Thermus thermophilus* (PDB ID: 2ZTJ, 28.9% identity, 45.5% similarity)^21^ was also aligned properly with HphA. These two HCSs were utilized to analyze the amino acid residues in the active site (Figure 2B). The amino acid sequence of HphA was also aligned with three structurally characterized IPMSs from *Neisseria meningitidis* serogroup B (PDB: 3RMJ, 28.7% identity, 44.9% similarity),^22^ from *Cytophaga hutchinsonii* (PDB: 3EEG, 26.2% identity, 40.5% similarity),^23^ and from *Listeria monocytogenes* serotype 4b str. F2365 (PDB ID: 3EWB, 28.5% identity, 44.6% similarity)^24^ (Figure 2C). These bioinformatic analyses identified five amino acid residues that may play a role in the substrate selection, which were A73, D157, A159, M186, and S242 in HphA (Table 1).

**Table 1.** The amino acid residues in the active site of HphA and homologous enzymes, which were hypothesized to play roles in the substrate selection.

| Enzymes | HphA | HCS | IPSM | hHphA |
| --- | --- | --- | --- | --- |
| **Amino acid residues** | Ala73 | His | Leu | *Ala* |
|  | Asp157 | Arg | Glu | Asn |
|  | Ala159 | Thr | Ser | *Ala* |
|  | Met186 | Ser | Asn | Phe |
|  | Ser242 | His | Glu | *Ser* |

The amino acid residues that are identical in the homologous enzymes to HphA are italicized. Each abbreviation depicts as follows: HCS = homocitrate synthase, IPMS = 2-isopropylmalate synthase, hHphA = homologous enzyme of HphA that was proposed to accept homologated 4-hydroxyphenylpyruvic acid as a substrate.

**Figure 2.**
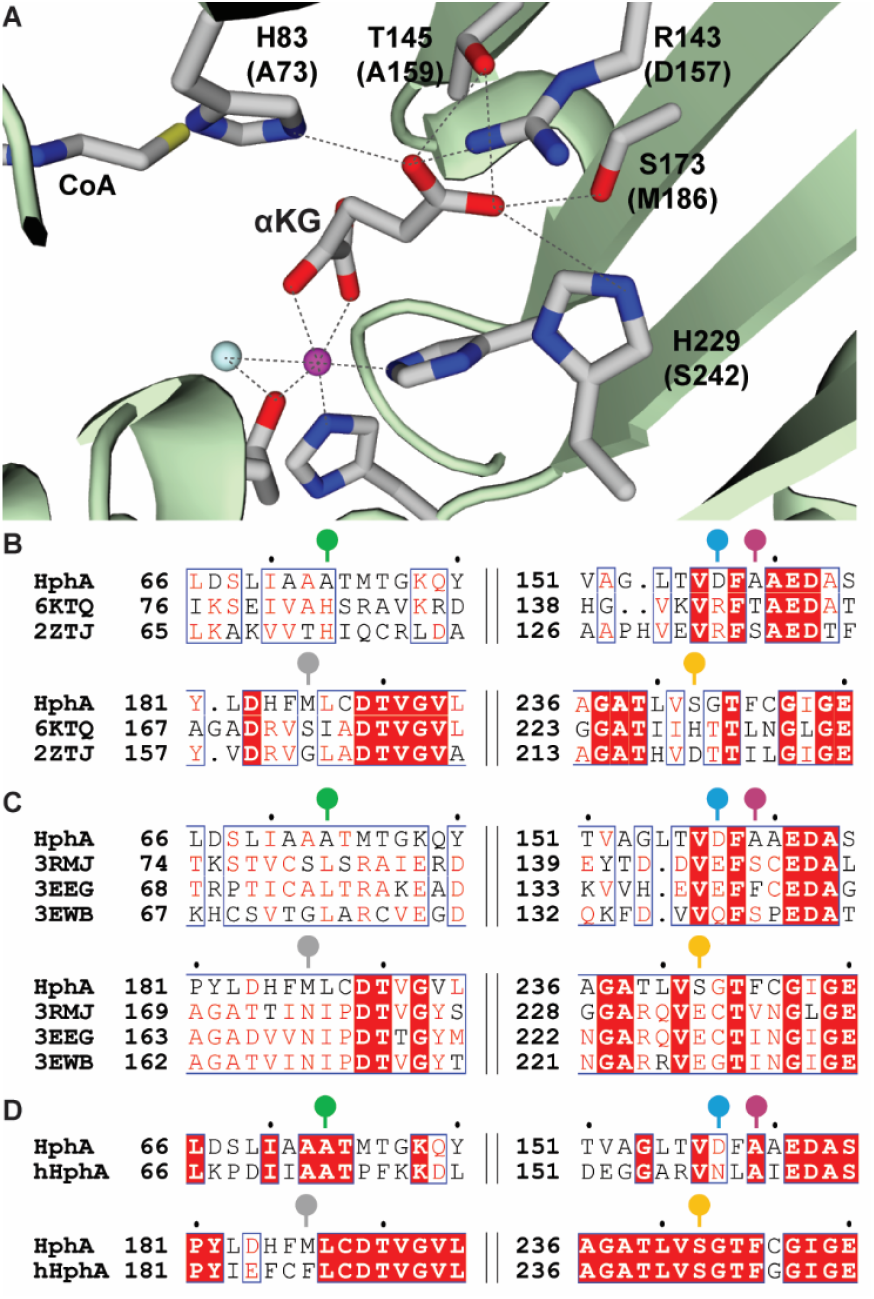
**A**. The active site of homocitrate synthase with αKG and CoA, focusing on interactions between the side chain of αKG and the amino acid residues of HCS (PDB ID: 6KTQ).^19^ The magenta and cyan balls are Zn^2+^ and a water molecule, respectively. The amino acid residues in parentheses are those found in HphA. The dotted lines represent potential polar interactions. The amino acid sequence alignments are shown as follows: **B**. HphA, HCS from *S. acidocaldarious* (PDB ID: 6KTQ),^19^ and HCS from *T. thermophilus* (PDB ID: 2ZTJ),^21, 22^ **C**. HphA, IPMS from *N. meningitidis* serogroup B (PDB ID: 3RMJ),^22^ IPMS from *C. hutchinsonii* (PDB: 3EEG),^23^ and IPMS from *L. monocytogenes* serotype 4b str. F2365 (PDB ID: 3EWB),^24^ and **D**. HphA and hHphA from *M. aeruginosa* NIES-4285.^14^ The amino acid residues targeted in this study are represented by sticks with circular heads. The colors used correspond to those in Figures 3B-E.

In this study, the amino acid sequence of recently-discovered hHphA from *M. aeruginosa* NIES-4285, which has not been characterized functionally or structurally, was additionally aligned with HphA (Figure 2D and Figure S1).^14^ HphA and hHphA are highly homologous and display 71.7% sequence identity and 82.7% sequence similarity to each other. Based on this sequence alignment and the previous consensus of HCS and IPMS data with HphA, the M186 residue was the only residue that was not conserved in HphA and hHphA and was therefore predicted to be the key residue for expanding the substrate promiscuity to a slightly larger side chain. Based on the bioinformatics data, we designed mutations to mimic the activity of HCS, IPMS, or hHphA; specific amino acid mutations are shown in Table 1.

### Substrate alteration of single mutants that mimic the homologous enzymes

The catalytic functions of the single mutants of HphA, which mimic the active site of the homologous enzymes HCS and IPMS, were evaluated through *in vitro* enzymatic assays. Their relative activities were measured through colorimetric assays, using the substrates αKG and OAA for the HCS mimic as well as MOBA for the IPMS mimic (Figure 3A). Since the reactions catalyzed by HphA and its homologous enzymes produce CoA as a co-product, Ellman’s reagent was used to detect the free thiol.^13^ Activity was determined after the reactions had proceeded for 20 minutes; each reactivity was related to HphA WT with the same substrate, which was set to be 100% activity (Figures 3B-E).

**Figure 3.**
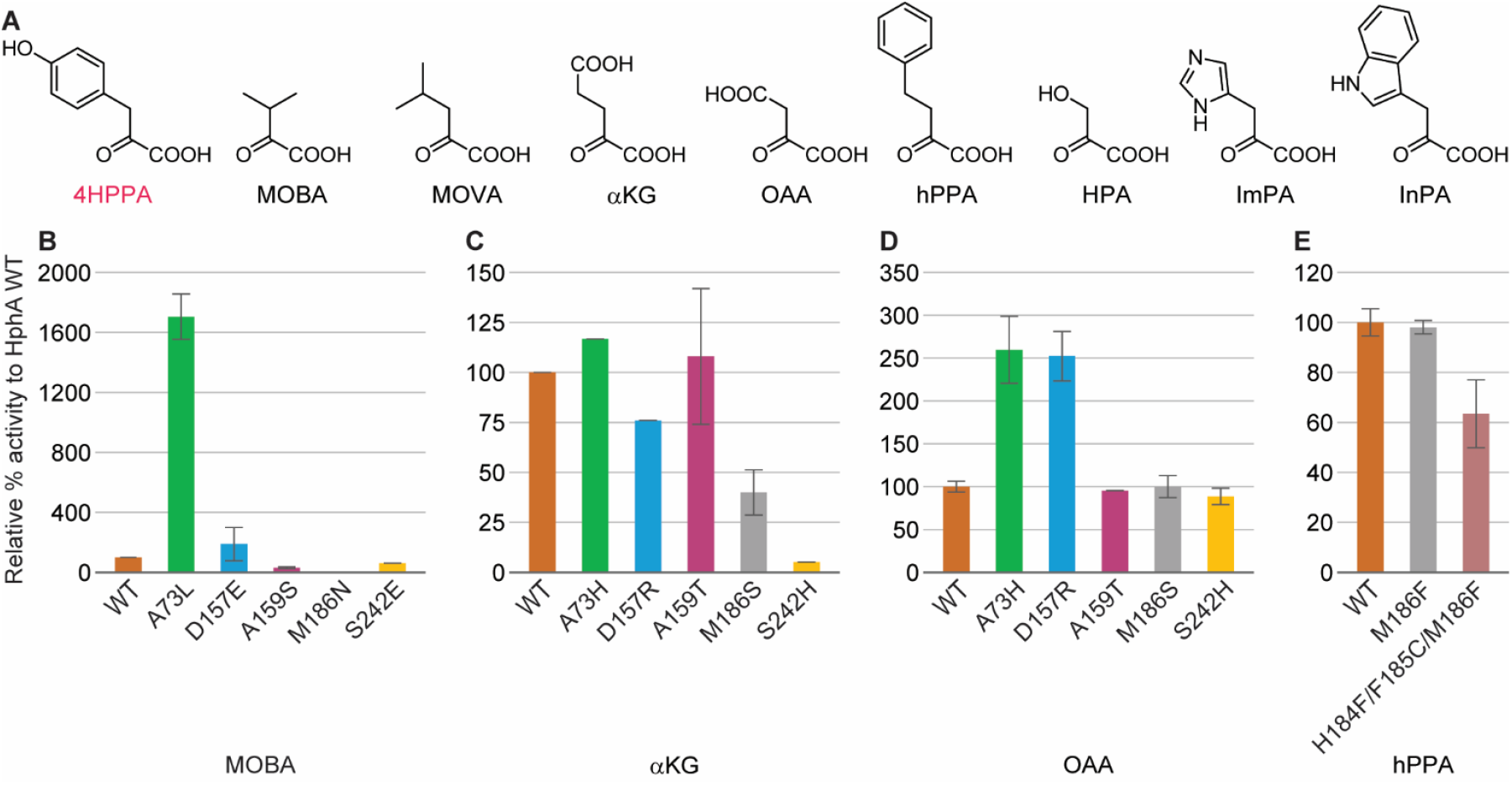
**A**. Structures of small molecules used in this study. **B-E**. Relative % activity of HphA mutants to HphA WT for mimicking **B**. the IPMS active site with MOBA, **C**. the HCS active site with αKG, **D**. the HCS active site with OAA, and **E**. the hHphA active site with hPPA. Each abbreviation depicts as follows: 4HPPA = 4-hydroxyphenylpyruvic acid, MOBA = 3-methyl-2-oxobutanoic acid, MOVA = 4-methyl-2-oxovaleric acid, αKG = α-ketoglutaric acid, OAA = oxalacetic acid, hPPA = homologated pyruvic acid (2-oxo-4-phenylbutanoic acid), HPA = hydroxypyruvic acid, ImPA = 4-imidazolepyruvic acid, InPA = Indole-3-pyruvic acid. The error bars are the range of the data (n = 2).

HphA single mutants that mimic the IPMS active site were A73L, D157E, A159S, M186N, and S242E. The data revealed that HphA A73L enhanced the activity with MOBA 17-fold compared to the WT. HphA D157E displayed slightly higher activity than the WT (1.9-fold), while the other IPMS mimics showed less activity than the WT (Figure 3B).

HphA single mutants that mimicked the HCS active site included A73H, D157R, A159T, M186S, and S242H. These mutants were tested with αKG, but none of the mutants showed enhanced activity with the substrate compared to the WT (Figure 3C). Consequently, we tested the activity of these mutants with an analogous compound, OAA (Figure 3D). OAA is the substrate of citrate synthase (CS), which is another homologous enzyme to HphA and is involved in the citric acid cycle in eukaryotes.^25^ OAA was approximately 2.5-fold better as a substrate for both HphA A73H and D157R as compared to the WT, while the other mutants did not display enhanced activity with this compound.

The data obtained from the single mutants designed to mimic the active sites of IPMS and HCS suggests that the A73 residue plays a role in substrate selectivity. These results were unexpected based on the bioinformatic analysis of the active site of hHphA, as discussed in the previous section (*Bioinformatics to design the mutant that mimics the activity of homologous enzymes*), predicting the importance of the M186 residue in the substrate selectivity of HphA. This was confirmed by evaluating the activity of HphA mutants that mimic the hHphA active site with hPPA, including the single mutant M186F and the triple mutant H184F/F185C/M186F which residues form a portion of the β-sheet in the active site. Neither of these mutants showed increased activity with hPPA (Figure 3E).

One of the IPMS mimics, A73L, displayed a significantly greater increase in activity with its substrate compared to the HCS mimic, A73H. This difference can be attributed to the distinct nature of the substrate side chains. Specifically, the types of substrates for each enzyme are as follows: HphA interacts with non-polar or slightly polar aromatic groups (Phe and Tyr), IPMS with non-polar aliphatic group (Val), and HCS with polar acidic group (Glu). Therefore, it is reasonable to conclude that mimicking the IPMS active site is simpler than that of the HCS active site. As a result, we expanded the set of HphA mutants designed to mimic the HCS active site by introducing additional mutations.

### Expansion of the mutations for polar substrates

To enhance the HCS mimic activity with αKG and OAA, we created double mutants of HphA based on the A73H mutation. We incorporated A73H into each of the double mutants, due to its apparent impact on substrate selectivity and demonstrated higher activity with OAA than HphA WT. The other mutation in each double mutant was selected from the previously tested single mutants. We particularly anticipated positive outcomes from the A73H/D157R combination, while still considering A159T, M186S, and S242H as potentially important residues in the active site.

The results of activity tests of the double mutants as compared to HphA WT and A73H are shown in Figure 4. Unfortunately, none of the double mutants had significantly more activity against OAA and/or αKG than the single A73H mutant; even the A73H/D157R double mutant did not have higher activity, contradicting expectations. It is important to note that all double mutations caused the enzyme to be less stable: reaction solutions began to form precipitate after 30 min of incubation at room temperature.

**Figure 4.**
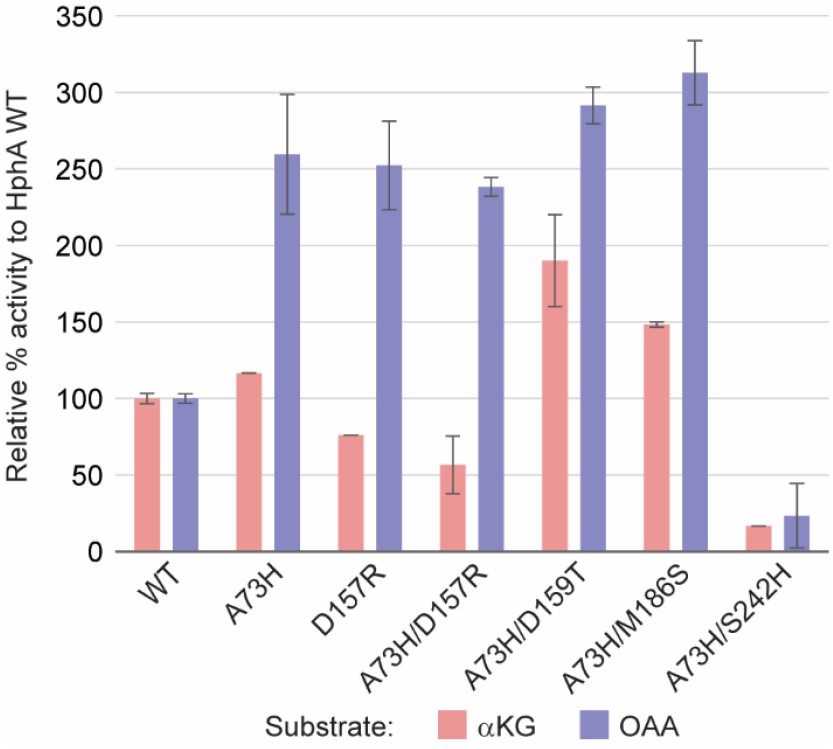
Relative % activity to HphA WT of HphA double mutants that mimic the HCS active site with αKG (pink) and OAA (blue) as substrates. The error bars are the range of the data (n = 2).

### Expansion of the A73 mutations for various substrates

Following evaluation of the single mutants designed to mimic the active sites of IPMS and HCS, the A73 residue was studied further to create mutants that accept a wide range of substrates. Other than A73L and A73H, we designed six more mutations from A73 and tested their activities compared to HphA WT on nine different substrates.

Based on the data obtained from HphA A73L and A73H, we selected to mutate the A73 residue to V, G, F, N, D, and S. The A73V mutant was designed to slightly expand the active site binding pocket from the A73L mutant, anticipating that MOBA and/or MOVA (containing the Leu side chain) could act as substrates. The A73G mutant was developed to increase activity with compounds larger than the natural substrates of HphA WT, phenylpyruvic acid (PPA) and 4HPPA, such as hPPA. Conversely, the A73F mutant was aimed at making the active site smaller to see if the substrate scope was modified accordingly. The mutations to increase the polarity, which were A73N, A73D, and A73S, were intended to create higher flexibility in the active site than the A73H mutation for the substrates with polar side chains.

Each mutation was tested with nine different commercially available substrates (Figures 3A and 5). 4HPPA is one of the natural substrates of HphA and was used as a control. MOBA, αKG, and OAA are the natural substrates of IPMS, HCS, and CS, respectively, as discussed in previous sections. MOVA is the homologated version of MOBA and was included to evaluate if the active site with a smaller binding pocket better utilizes this compound. hPPA was used to test if di-homologation can be initiated by a mutant HphA. HPA is the Ser counterpart of PPA and 4HPPA. All the above homologated compounds can be found in nature. InPA and ImPA are the Trp and His counterparts of PPA and 4HPPA, respectively, which homologated versions (hTrp and hHis) have not been found in nature.

The activity of the mutants with each substrate was compared to the activity of HphA WT with 4HPPA (Figure 5). Contrary to our expectations, HphA A73L, which was intended to mimic the IPMS active site, was the most active of all the mutants across all substrates, except for hPPA, ImPA, and αKG. The A73L mutant showed activity with 4HPPA (133.0%), InPA (5.3%), MOBA (85.6%; as also presented in Figure 3B), MOVA (73.9%), HPA (6.4%), and OAA (7.1%). All mutations showed negligible activity with ImPA or αKG, while hPPA was only slightly active with the A73G mutant (2.0%). The polar substrates, ImPA, HPA, OAA, and αKG, were anticipated to be more active with the polar mutations (A73N, A73D, A73S, and A73H), which was not supported by the data. Mutations to large amino acid side chains, A73F and A73H, almost completely disrupted the HphA activity, including that with 4HPPA. These results may not be only because of the disruption of the substrate binding; since the A73 residue is also close to the CoA binding site (Figure 2A), mutations to the large side chains may damage the binding of CoA in the active site.

**Figure 5.**
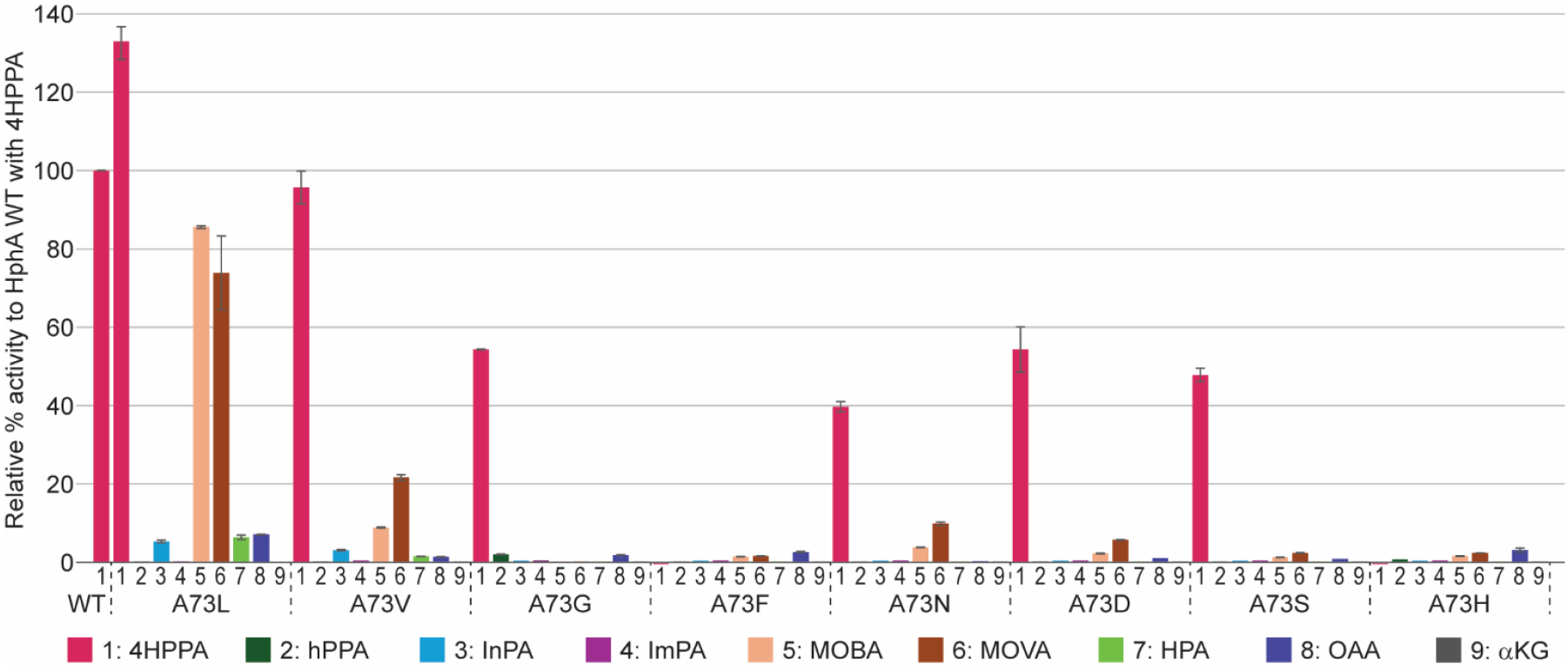
Relative % activity of A73 series mutant with various substrates to HphA WT with the natural substrate 4HPPA. The error bars are the range of the data (n = 2).

### Kinetics of HphA A73L compared to homologous enzymes

To better understand the reactions catalyzed by HphA A73L, the Michaelis-Menten kinetic parameters were determined with active substrates, including 4HPPA, InPA, MOBA, MOVA, HPA, and OAA. Michaelis-Menten parameters were also determined for HphA WT with MOBA and MOVA, IPMS with MOBA and MOVA, as well as HCS with αKG and OAA (Table 2). Kinetic data of HphA WT with 4HPPA were reported previously, with a *k*_cat_/*K*_m_ of 3.4 ± 0.37 mM^-1^ s^-1^.^12^ All kinetic assays were performed using the same methods as those for HphA with 4HPPA, except slight modifications made for HCS. Due to the instability of HCS under the assay conditions, its stability was enhanced by adding 10% glycerol and starting the reactions on ice. As a result, the comparisons of activity between HphA A73L and HCS were not very reliable.

**Table 2.** Kinetic data for HphA WT, homologous enzymes, and HphA A73L mutant.

| Enzyme | Substrate | $k_{\text{cat}}$ ($\text{s}^{-1}$) | $K_{\text{m}}$ (mM) | $k_{\text{cat}}/K_{\text{m}}$ ($\text{mM}^{-1} \text{s}^{-1}$) |
| --- | --- | --- | --- | --- |
| **HphA WT** | 4HPPA<sup>12</sup> | $0.14 \pm 0.0025$ | $0.040 \pm 0.0042$ | $3.4 \pm 0.37$ |
| | MOBA | $0.0078 \pm 0.0008$ | $4.0 \pm 1.1$ | $0.0019 \pm 0.0006$ |
| | MOVA | $0.032 \pm 0.008$ | $7.0 \pm 2.7$ | $0.0046 \pm 0.0021$ |
| **IPMS** | MOBA | $0.70 \pm 0.05$ | $0.11 \pm 0.04$ | $6.4 \pm 2.1$ |
| | MOVA | $0.013 \pm 0.001$ | $0.011 \pm 0.002$ | $1.1 \pm 0.2$ |
| **HCS** | OAA | $0.17 \pm 0.23$ | $24.0 \pm 38.5$ | $0.0071 \pm 0.0143$ |
| | αKG | $0.065 \pm 0.005$ | $0.92 \pm 0.15$ | $0.070 \pm 0.013$ |
| **HphA A73L** | 4HPPA | $0.22 \pm 0.01$ | $0.41 \pm 0.05$ | $0.54 \pm 0.08$ |
| | MOBA | $0.12 \pm 0.01$ | $0.65 \pm 0.15$ | $0.18 \pm 0.05$ |
| | MOVA | $0.10 \pm 0.01$ | $1.63 \pm 0.35$ | $0.064 \pm 0.015$ |
| | OAA | $0.0069 \pm 0.0002$ | $0.0030 \pm 0.0006$ | $2.3 \pm 0.5$ |
| | HPA | $0.0032 \pm 0.0005$ | $0.30 \pm 0.15$ | $0.011 \pm 0.006$ |
| | InPA | $0.046 \pm 0.014$ | $1.8 \pm 1.2$ | $0.025 \pm 0.018$ |

4HPPA = 4-hydroxyphenylpyruvic acid; MOBA = 3-methyl-2-oxobutanoic acid; MOVA = 4-methyl-2-oxovaleric acid; OAA = 2-oxosuccinic acid; HPA = hydroxypyruvic acid; InPA = indole-3-pyruvic acid; αKG = $\alpha$ -ketoglutarate. The data for HphA WT with 4HPPA was obtained in a previous study.<sup>12</sup>

The kinetics of HphA A73L with 4HPPA showed a *k*_cat_/*K*_m_ of 0.54 ± 0.08 mM^-1^ s^-1^, which was 6.3-fold lower than that of HphA WT. This result was anticipated because the A73L mutation was designed to accept IPMS substrates. The kinetics for IPMS with MOBA and MOVA showed that *k*_cat_/*K*_m_ values of 6.4 ± 2.1 mM^-1^ s^-1^ and 1.1 ± 0.2 mM^-1^ s^-1^, respectively. IPMS was very efficient with MOBA, its natural substrate, as well as decently efficient with MOVA, the homologated version of MOBA. As compared to these values, HphA WT displayed *k*_cat_/*K*_m_ values of 3300-fold and 240-fold lower for MOBA and MOVA, respectively. The poor activity of HphA WT with MOBA and MOVA was previously demonstrated qualitatively, as MOBA showed only a few % activity compared to 4HPPA.^12^ The *k*_cat_/*K*_m_ values for HphA A73L with MOBA and MOVA were anticipated to be greater than those of HphA WT due to its mutation to be an IPMS mimic. Its *k*_cat_/*K*_m_ value for MOBA was 0.18 ± 0.05 mM^-1^ s^-1^, which was 35.3-fold lower and 95-fold higher than those of IPMS and HphA WT, respectively. Similarly, the HphA A73L kinetic value for MOVA was 0.064 ± 0.015 mM^-1^ s^-1^, which was 17-fold lower and 14-fold higher than those of IPMS and HphA WT, respectively. These values showed that the A73L mutation on HphA significantly expanded the substrate scope for MOBA and MOVA, but it still does not reach the activity level of IPMS.

The kinetics of HphA A73L with OAA yielded a *k*_cat_/*K*_m_ value of 2.3 ± 0.5 mM^-1^ s^-1^, which was 318-fold higher than the *k*_cat_/*K*_m_ of HCS with OAA (0.0071 ± 0.0143 mM^-1^ s^-1^). This was likely because OAA is the natural substrate of CS, although it is homologous to HCS. The high error associated with HCS and OAA was due to the high *K*_m_ value and the inhibition of the enzyme activity with a substrate concentration higher than 5 mM. Since the A73L mutation was intended to mimic the IPMS activity, rather than that of HCS, it was previously anticipated that this mutant would have no or minimum activity with OAA. However, a high *k*_cat_/*K*_m_ value for HphA A73L with OAA was observed due to a significantly low *K*_m_ value of 0.0030 ± 0.0006 mM compared to other substrates, although it displayed a very low *k*_cat_ value of 0.0069 ± 0.0002 s^-1^. These data suggest that the interactions between HphA A73L and OAA are very strong, but the product is not efficiently released from the enzyme’s active site. This was also observed in the low production of CoA in the substrate profile experiments involving HphA A73L (Figure 5).

Kinetics of HphA A73L with HPA and InPA were also obtained; the *k*_cat_/*K*_m_ of this mutant with HPA was 0.011 ± 0.006 mM^-1^ s^-1^. Since HPA contains an alcohol group on its side chain, unlike MOBA which HphA A73L was originally designed to accept, it was not anticipated that the A73L mutant would accept this compound as a substrate. However, HphA A73L retained some activity with HPA; its activity was 6.1% and 17% to that of HphA A73L with MOBA and MOVA, respectively. This slight activity could be due to its similar size of the side chain to MOBA. The *k*_cat_/*K*_m_ value for HphA A73L with InPA was 0.025 ± 0.018 mM^-1^ s^-1^. Although it was the lowest among the active substrates for this mutant, it was measurable and significantly higher than that of HphA WT with MOBA and MOVA. These results demonstrated that HphA A73L has a wider range of substrate scopes than we anticipated from the bioinformatic study, showing that HphA is amenable to engineering in its active site.

## CONCLUSION

HphA, which is part of the homologation pathway in secondary metabolites of cyanobacteria, has previously been shown to be a substrate-specific enzyme and acts as the gatekeeper of the pathway. The homologation pathway, once characterized and engineered, has the potential to be used in derivatization of bioactive peptide molecules, so the goal of this project was to demonstrate the amenability of HphA to engineering for future research. Through bioinformatic studies using the homologous enzymes HCS, IPMS, and hHphA, several potential amino acid residues in the active sites were revealed for the alteration of the substrate scope; these locations were targeted for mutations of HphA. Time-course assays with the mutants, mimicking the active sites of HCS and IPMS, identified the A73 residue as key in the substrate specificity of HphA. Subsequently, a total of eight mutations on the A73 residue were conducted to expand or alter the substrate scope. The A73L mutant demonstrated activity with a much wider range of substrates than the WT; these substrates included amino acid counterparts of Val, Leu, Ser, Trp, and Asp in addition to the natural substrate of the WT, Tyr, out of nine substrates tested. Significantly, activity of this mutant with the Val counterpart, MOBA, increased 95-fold from that of the WT. Moreover, acceptance of the Trp counterpart, InPA, as a substrate was intriguing, as the homologated Trp (hTrp) does not occurr in nature, and the synthetic compound is not readily commercially available. Optimizing the substrate scope for Trp would lead to the production of hTrp by the homologation pathway, which will provide an additional scaffold for drug synthesis. As only one mutation in the bioinformatically analyzed active site significantly relaxed the substrate specificity, it could be optimized further for specific noncanonical substrates or for higher promiscuity by other methods, such as structure-guided mutagenesis following X-ray crystallography and directed evolution. This initial engineering study to prove the amenability of HphA promises hopeful applications of the homologation pathway.

## ASSOCIATED CONTENT

## Data Availability Statement

The data that support the findings of this study are available in the supporting information (SI) of this article in a PDF format. SI Figure include the full amino acid sequencing alignment between HphA and hHphA (Figure S1); Michaelis-Menten kinetic plots of HphA A73L (Figure S2); Michaelis-Menten kinetic plots of HphA WT, IPMS, and HCS (Figure S3); and PCR primers used in this study (Table S1).

## ACCESSION CODES

HphA: NCBI WP_012409019; UniProt B2J8P8

hHphA; NCBI WP_130757515; UniProt A0A402DEV0

HCS from *S. acidocaldarious*: PDB 6KTQ; UniProt Q4J989

HCS from *T. thermophilus*: PDB 2ZTJ; UniProt O87198

IPMS from *N. meningitidis* serogroup B: PDB 3RMJ; UniProt Q9JZG1

IPMS from *C. hutchinsonii*: PDB 3EEG; UniProt A0A7D9NMJ7

IPMS from *L. monocytogenes* serotype 4b str. F2365: PDB 3EWB; UniProt Q71Y35

## Author Contributions

Study design: S.M. and R.M.L.H.; manuscript preparation: S.M., R.M.L.H., G.B., and A.C.S.; data generation and analysis: S.M., R.M.L.H., G.B., F.O.A., S.R.P., M.S., R.K.N., D.B.D., and A.C.S.; CHEM 4533 supervision: A.C.S.; funding acquisition: S.M.

## Notes

The authors declare no conflict of interest.

## ACKNOWLEDGEMENT

The preparation of this publication was supported by the following funding sources: National Institute of General Medical Sciences of the National Institutes of Health under award number 1R15GM151721-01, start-up funds from the Department of Chemistry and Biochemistry at Augusta University, and the Summer Scholars Program as well as Materials Grants in the Center for Undergraduate Research and Scholarship at Augusta University. We would like to thank CHEM 4533 students at Augusta University in Spring 2025, who created several HphA mutants contributing to this study. This course delivery was supported by the Department of Chemistry and Biochemistry at Augusta University.

## Notes

### Competing Interest Statement

The authors have declared no competing interest.

## REFERENCES

1. Les, F.; Valero, M. S.; Arruebo, M. P., Natural Product Chemistry and Biological Research. Int J Mol Sci 2024, 25 (7).

2. Atanasov, A. G.; Zotchev, S. B.; Dirsch, V. M.; International Natural Product Sciences, T.; Supuran, C. T., Natural products in drug discovery: advances and opportunities. Nat Rev Drug Discov 2021, 20 (3), 200–216.

3. Newman, D. J.; Cragg, G. M., Natural Products as Sources of New Drugs over the Nearly Four Decades from 01/1981 to 09/2019. J Nat Prod 2020, 83 (3), 770–803.

4. Wenski, S. L.; Thiengmag, S.; Helfrich, E. J. N., Complex peptide natural products: Biosynthetic principles, challenges and opportunities for pathway engineering. Synth Syst Biotechnol 2022, 7 (1), 631–647.

5. McIntosh, J. A.; Donia, M. S.; Schmidt, E. W., Ribosomal peptide natural products: bridging the ribosomal and nonribosomal worlds. Nat Prod Rep 2009, 26 (4), 537–59.

6. Ding, Y.; Ting, J. P.; Liu, J.; Al-Azzam, S.; Pandya, P.; Afshar, S., Impact of nonproteinogenic amino acids in the discovery and development of peptide therapeutics. Amino Acids 2020, 52 (9), 1207–1226.

7. Owens, S. L.; Ahmed, S. R.; Lang Harman, R. M.; Stewart, L. E.; Mori, S., Natural Products That Contain Higher Homologated Amino Acids. ChemBioChem 2024, 25 (9), e202300822.

8. Monteiro, P. R.; do Amaral, S. C.; Siqueira, A. S.; Xavier, L. P.; Santos, A. V., Anabaenopeptins: What We Know So Far. Toxins (Basel) 2021, 13 (8).

9. Koketsu, K.; Mitsuhashi, S.; Tabata, K., Identification of homophenylalanine biosynthetic genes from the cyanobacterium Nostoc punctiforme PCC73102 and application to its microbial production by Escherichia coli. Appl Environ Microbiol 2013, 79 (7), 2201–8.

10. Zhang, K.; Sawaya, M. R.; Eisenberg, D. S.; Liao, J. C., Expanding metabolism for biosynthesis of nonnatural alcohols. Proc Natl Acad Sci U S A 2008, 105 (52), 20653–8.

11. Petersen, A.; Hansen, L. G.; Mirza, N.; Crocoll, C.; Mirza, O.; Halkier, B. A., Changing substrate specificity and iteration of amino acid chain elongation in glucosinolate biosynthesis through targeted mutagenesis of Arabidopsis methylthioalkylmalate synthase 1. Biosci Rep 2019, 39 (7).

12. Stewart, L. E.; Owens, S. L.; Ahmed, S. R.; Lang Harman, R. M.; Mori, S., Characterization of HphA: The First Enzyme in the Homologation Pathway of L-Phenylalanine and L-Tyrosine. ChemBioChem 2024, 25 (16), e202400369.

13. Ellman, G. L., Tissue sulfhydryl groups. Arch Biochem Biophys 1959, 82 (1), 70–7.

14. Phan, C. S.; Ling, Z.; Mehjabin, J. J.; Matsuda, K.; Prakoso, N. I.; Umezawa, T.; Wakimoto, T.; Okino, T., Doubly Homologated Tyrosine-Containing Peptides from the Cyanobacterium Microcystis aeruginosa NIES-4285 and Their Biosynthesis. J Nat Prod 2024, 87 (11), 2629–2639.

15. Corpet, F., Multiple sequence alignment with hierarchical clustering. Nucleic Acids Res 1988, 16 (22), 10881–90.

16. Higuchi, R.; Krummel, B.; Saiki, R. K., A general method of in vitro preparation and specific mutagenesis of DNA fragments: study of protein and DNA interactions. Nucleic Acids Res 1988, 16 (15), 7351–67.

17. de Carvalho, L. P.; Blanchard, J. S., Kinetic and chemical mechanism of alpha-isopropylmalate synthase from Mycobacterium tuberculosis. Biochemistry 2006, 45 (29), 8988–99.

18. Kumar, V. P.; West, A. H.; Cook, P. F., Kinetic and chemical mechanisms of homocitrate synthase from Thermus thermophilus. J Biol Chem 2011, 286 (33), 29428–29439.

19. Suzuki, T.; Tomita, T.; Hirayama, K.; Suzuki, M.; Kuzuyama, T.; Nishiyama, M., Involvement of subdomain II in the recognition of acetyl-CoA revealed by the crystal structure of homocitrate synthase from Sulfolobus acidocaldarius. FEBS J 2021, 288 (6), 1975–1988.

20. Zhang, Z.; Wu, J.; Lin, W.; Wang, J.; Yan, H.; Zhao, W.; Ma, J.; Ding, J.; Zhang, P.; Zhao, G. P., Subdomain II of alpha-isopropylmalate synthase is essential for activity: inferring a mechanism of feedback inhibition. J Biol Chem 2014, 289 (40), 27966–78.

21. Okada, T.; Tomita, T.; Wulandari, A. P.; Kuzuyama, T.; Nishiyama, M., Mechanism of substrate recognition and insight into feedback inhibition of homocitrate synthase from Thermus thermophilus. J Biol Chem 2010, 285 (6), 4195–4205.

22. Huisman, F. H.; Koon, N.; Bulloch, E. M.; Baker, H. M.; Baker, E. N.; Squire, C. J.; Parker, E. J., Removal of the C-terminal regulatory domain of alpha-isopropylmalate synthase disrupts functional substrate binding. Biochemistry 2012, 51 (11), 2289–97.

23. Sugadev, R.; Burley, S. K.; Swaminathan, S., Crystal structure of a 2-isopropylmalate synthase from Cytophaga hutchinsonii. 2008, 10.2210/pdb3EEG/pdb.

24. Ramagopal, U. A.; Toro, R.; Gilmore, M.; Hu, S.; Maletic, M.; Rodgers, L.; Burley, S. K.; Almo, S. C., Crystal structure of N-terminal domain of putative 2-isopropylmalate synthase from Listeria monocytogenes. 2008, 10.2210/pdb3EWB/pdb.

25. Chhimpa, N.; Singh, N.; Puri, N.; Kayath, H. P., The Novel Role of Mitochondrial Citrate Synthase and Citrate in the Pathophysiology of Alzheimer’s Disease. J Alzheimers Dis 2023, 94 (1), S453–S472.

