## Supplementary material for "Amenability to Engineering of the Homologation Enzyme, HphA, through Homologous-Based Site-Directed Mutagenesis": Figure S1; Figure S2; Figure S3; Table S1

**Outline:**

**Figures and Tables**

Figure S1. Amino acid sequence alignment between HphA and hHphA 3

Figure S2. Kinetic assays 4

Figure S3. Kinetic assays 2 5

Table S1. PCR primers used in this study 6

**References** 7

### 2. Figures and Tables

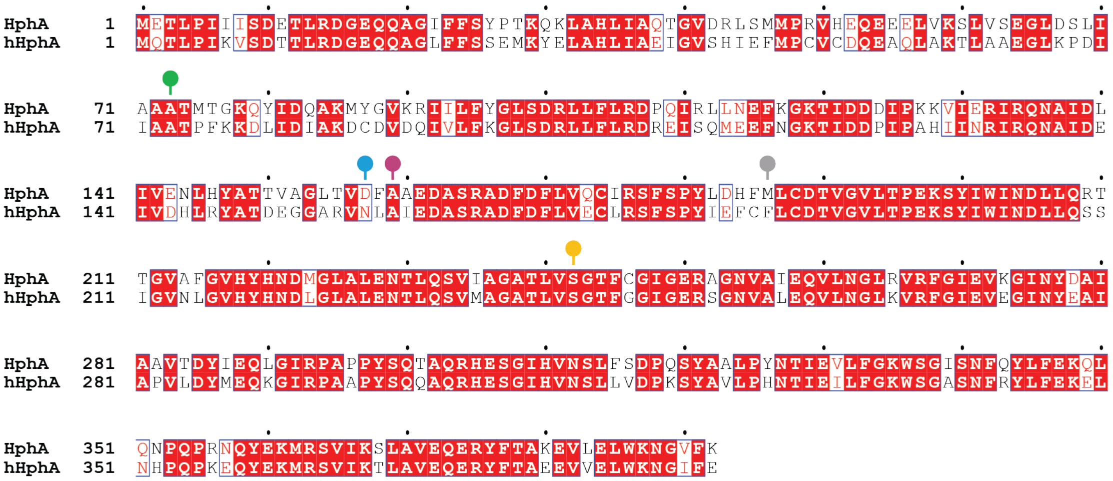


**Figure S1.** The full amino acid sequencing alignment between HphA and hHphA.[1, 2] Each pin represents the potential amino acid residue in the active site, which play a role in the substrate selectivity.

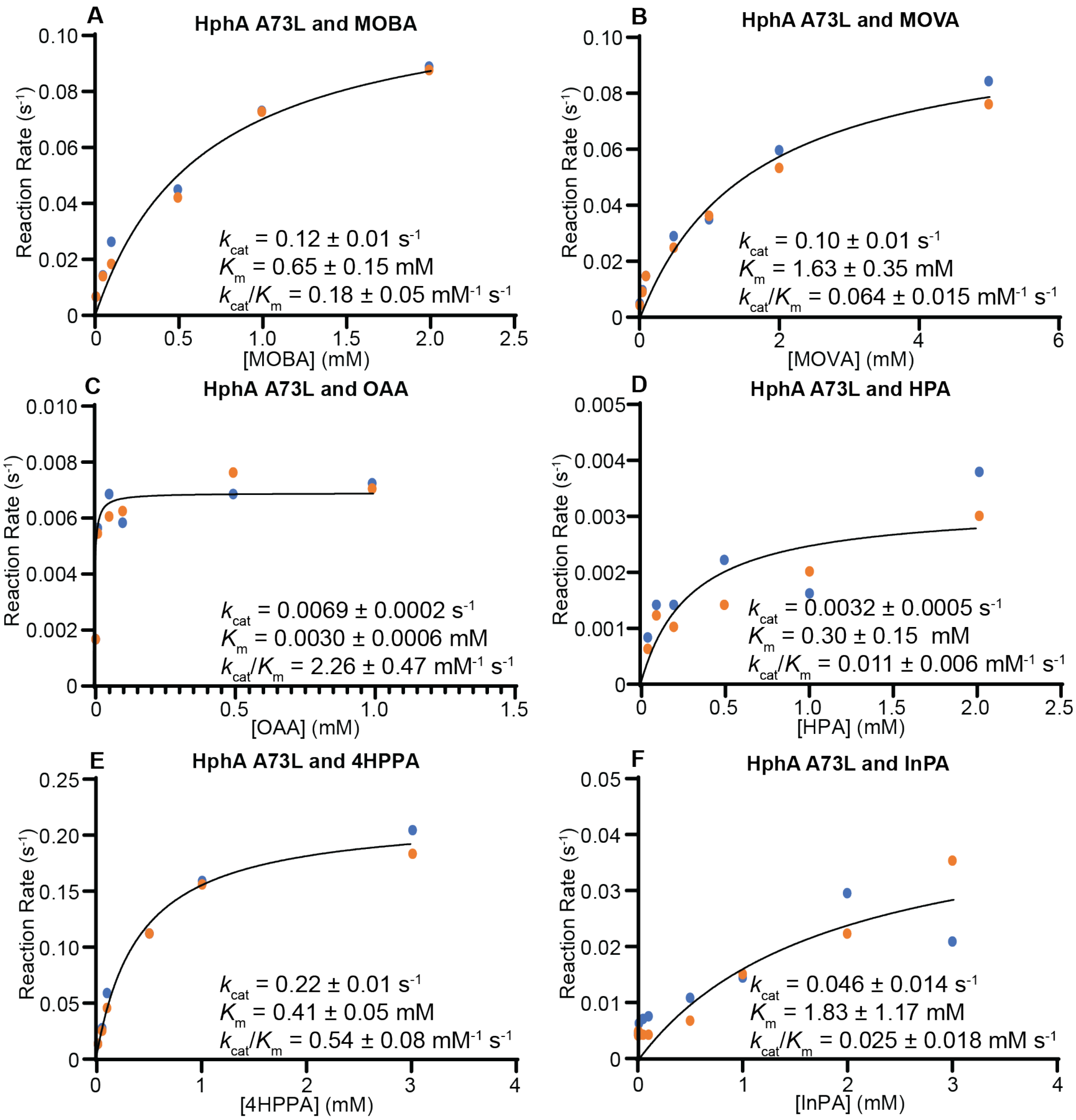


**Figure S2.** Michaelis–Menten kinetics of HphA A73L with various substrates. **A.** HphA A73L with 3-methyl-2-oxobutanoic acid (MOBA); **B.** HphA A73L with 4-methyl-2-oxovaleric acid (MOVA); **C.** HPhA A73L with oxaloacetic acid (OAA); **D.** HphA A73L with hydroxypyruvic acid (HPA); **E.** HphA A73L with 4- hydroxylphenylpyruvic acid (4HPPA); **F.** HphA A73L with indole-3-pyruvic acid (InPA). All assays were duplicated, and the data was individually plotted.

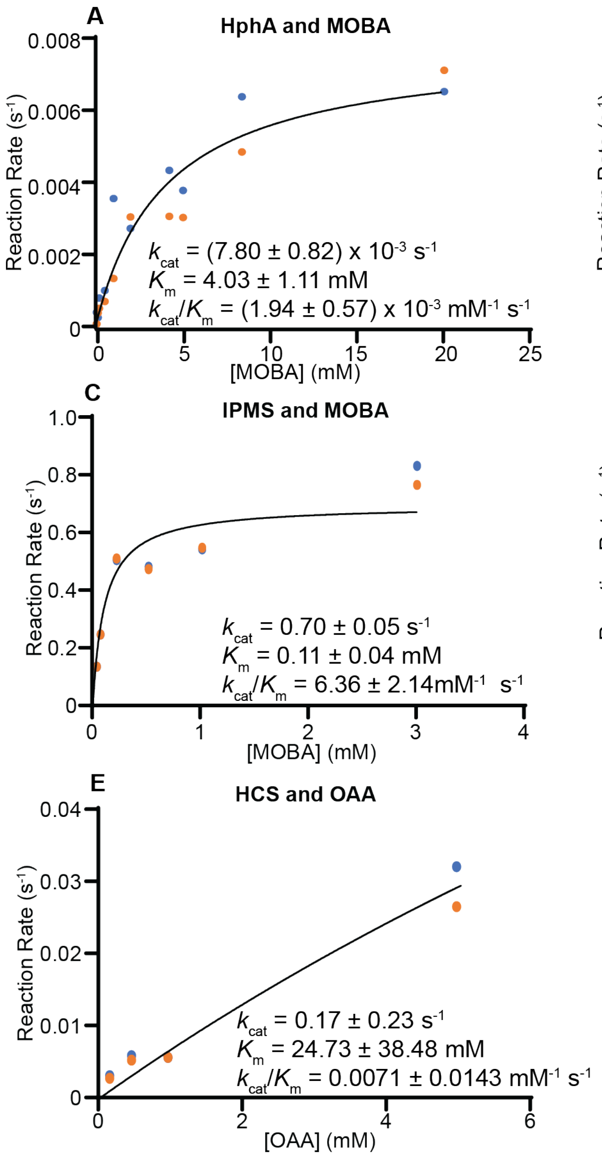


**Figure S3.** Michaelis–Menten Kinetics of HphA and homologous enzymes. **A.** HphA and 3-methyl-2-oxobutanoic acid (MOBA); **B.** HphA and 4-methyl-2-oxovaleric acid (MOVA); **C.** isopropylmalate synthase (IPMS) with MOBA; **D.** IPMS with MOVA; **E.** homocitrate synthase (HCS) with oxaloacetic acid (OAA); **F.** HCS with α-ketoglutaric acid (αKG). All assays were duplicated, and the data was individually plotted.

**Table S1.** PCR primers used in this study

| Primer # | Primer name | Primer sequence | 5' or 3' |
|---|---|---|---|
| 1 | HphA-NcoI-F | CTAAAACCATGGAAACCCTTCCTATCATA | 5' |
| 2 | HphA-XhoI-R-noSTOP | GAAGATCTCGAGTTTAAACACCCCATTTTTC | 3' |
| 3 | HphA-A73L-F | CTGATCGCTGCT***CTT***ACCATGACGGGA | 5' |
| 4 | HphA-A73L-R | TCCCGTCATGGT***AAG***AGCAGCGATCAG | 3' |
| 5 | HphA-D157E-F | GGACTCACTGTA***GAG***TTTGCCGCCGAAG | 5' |
| 6 | HphA-D157E-R | CTTCGGCGGCAAA***CTC***TACAGTGAGTCC | 3' |
| 7 | HphA-A159S-F | CACTGTAGACTTT***TCC***GCCGAAGATGCC | 5' |
| 8 | HphA-A159S-R | GGCATCTTCGGC***GGA***AAAGTCTACAGTG | 3' |
| 9 | HphA-M186N-F | CTCGACCATTTT***AAC***CTCTGTGATACC | 5' |
| 10 | HphA-M186N-R | GGTATCACAGAG***GTT***AAAATGGTCGAG | 3' |
| 11 | HphA-S242E-F | GCGACTTTAGTA***GAG***GGTACGTTTTGC | 5' |
| 12 | HphA-S242E-R | GCAAAACGTACC***CTC***TACTAAAGTCGC | 3' |
| 13 | HphA-A73H-F | CTGATCGCTGCT***CAT***ACCATGACGGGA | 5' |
| 14 | HphA-A73H-R | TCCCGTCATGGT***ATG***AGCAGCGATCAG | 3' |
| 15 | HphA-D157R-F | GGACTCACTGTA***CGC***TTTGCCGCCGAAG | 5' |
| 16 | HphA-D157R-R | CTTCGGCGGCAAA***GCG***TACAGTGAGTCC | 3' |
| 17 | HphA-A159T-F | CACTGTAGACTTT***ACC***GCCGAAGATGCC | 5' |
| 18 | HphA-A159T-R | GGCATCTTCGGC***GGT***AAAGTCTACAGTG | 3' |
| 19 | HphA-M186S-F | CTCGACCATTTT***AGC***CTCTGTGATACC | 5' |
| 20 | HphA-M186S-R | GGTATCACAGAG***GCT***AAAATGGTCGAG | 3' |
| 21 | HphA-S242H-F | GCGACTTTAGTA***CAC***GGTACGTTTTGC | 5' |
| 22 | HphA-S242H-R | GCAAAACGTACC***GTG***TACTAAAGTCGC | 3' |
| 23 | HphA-A73G-F | CTGATCGCTGCT***GGT***ACCATGACGGGA | 5' |
| 24 | HphA-A73G-R | TCCCGTCATGGT***ACC***AGCAGCGATCAG | 3' |
| 25 | HphA-A73V-F | CTGATCGCTGCT***GTT***ACCATGACGGGA | 5' |
| 26 | HphA-A73V-R | TCCCGTCATGGT***AAC***AGCAGCGATCAG | 3' |
| 27 | HphA-A73F-F | CTGATCGCTGCT***TTT***ACCATGACGGGA | 5' |
| 28 | HphA-A73F-R | TCCCGTCATGGT***AAA***AGCAGCGATCAG | 3' |
| 29 | HphA-A73S-F | CTGATCGCTGCT***TCT***ACCATGACGGGA | 5' |
| 30 | HphA-A73S-R | TCCCGTCATGGT***AGA***AGCAGCGATCAG | 3' |
| 31 | HphA-A73D-F | CTGATCGCTGCT***GAT***ACCATGACGGGA | 5' |
| 32 | HphA-A73D-R | TCCCGTCATGGT***ATC***AGCAGCGATCAG | 3' |
| 33 | HphA-A73N-F | CTGATCGCTGCT***AAT***ACCATGACGGGA | 5' |
| 34 | HphA-A73N-R | TCCCGTCATGGT***ATT***AGCAGCGATCAG | 3' |
| 35 | HphA-M186F-F | CTCGACCATTTT***TTC***CTCTGTGATACC | 5' |
| 36 | HphA-M186F-R | GGTATCACAGAG***GAA***AAAATGGTCGAG | 3' |
| 37 | HphA-H184F-F185C-M186F-F | CCTTATCTCGAC***TTTTGTTTC***CTCTGTGATACC | 5' |
| 38 | HphA-H184F-F185C-M186F-R | GGTATCACAGAG***GAAACAAAA***GTCGAGATAAGG | 3' |

*Note*: The underlined nucleotide sequences are the target of the corresponding restriction enzyme. The bold-italicized nucleotides are the mutation site. Primers #1 and #2 were designed in the previous study on HphA characterization.[3]

### References


1. Phan, C. S.; Ling, Z.; Mehjabin, J. J.; Matsuda, K.; Prakoso, N. I.; Umezawa, T.; Wakimoto, T.; Okino, T., Doubly Homologated Tyrosine-Containing Peptides from the Cyanobacterium Microcystis aeruginosa NIES-4285 and Their Biosynthesis. *J Nat Prod* **2024,** *87* (11), 2629-2639.
2. Corpet, F., Multiple sequence alignment with hierarchical clustering. *Nucleic Acids Res* **1988,** *16* (22), 10881-90.
3. Stewart, L. E.; Owens, S. L.; Ahmed, S. R.; Lang Harman, R. M.; Mori, S., Characterization of HphA: The First Enzyme in the Homologation Pathway of l-Phenylalanine and l-Tyrosine. *Chembiochem* **2024,** *25* (16), e202400369.
